# In-Pero: Exploiting deep learning embeddings of protein sequences to predict the localisation of peroxisomal proteins

**DOI:** 10.1101/2021.01.18.427146

**Authors:** Marco Anteghini, Vitor AP Martins dos Santos, Edoardo Saccenti

## Abstract

Peroxisomes are ubiquitous membrane-bound organelles, and aberrant localisation of peroxisomal proteins contributes to the pathogenesis of several disorders. Many computational methods focus on assigning protein sequences to subcellular compartments, but there are no specific tools tailored for the sub-localisation (matrix vs membrane) of peroxisome proteins. We present here In-Pero, a new method for predicting protein sub-peroxisomal cellular localisation. In-Pero combines standard machine learning approaches with recently proposed multi-dimensional deep-learning representations of the protein amino-acid sequence. It showed a classification accuracy above 0.9 in predicting peroxisomal matrix and membrane proteins. The method is trained and tested using a double cross-validation approach on a curated data set comprising 160 peroxisomal proteins with experimental evidence for sub-peroxisomal localisation. We further show that the proposed approach can be easily adapted (In-Mito) to the prediction of mitochondrial protein localisation obtaining performances for certain classes of proteins (matrix and inner-membrane) superior to existing tools. All data sets and codes are available at https://github.com/MarcoAnteghini and at www.systemsbiology.nl

## 1 Introduction

Eukaryotic cells are believed to be able to synthesise up to 10,000 different kinds of proteins, which evolved to function optimally in a specific subcellular localisation [1].

In eukaryotes, there are ten main subcellular localisations which can be further subdivided into intra-organellar compartments (Figure 1). These organelles perform one or more and often complementary specific tasks in the cellular machinery. Examples of organelles are the nucleus, for the storage of genetic (DNA) material, mitochondria for the production of energy and the peroxisome, oxidative organelles.

**Figure 1.**
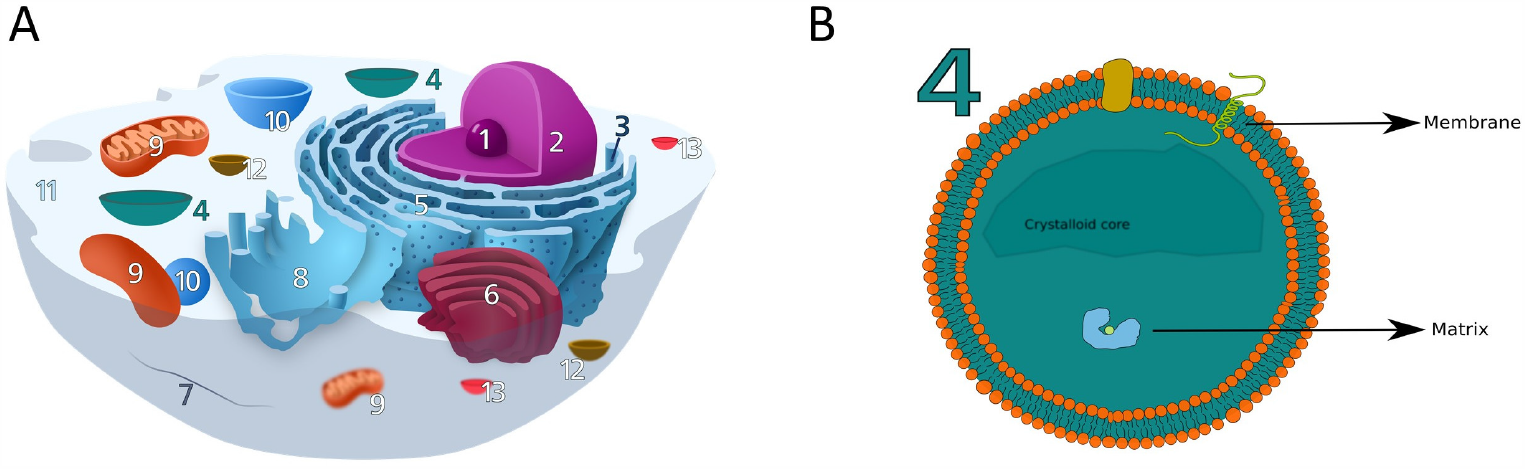
A) The eukaryotic cell and its organelles and compartments: (1) Nucleolus, Nucleus, (3) Ribosome, (4) Peroxisome, (5) Rough Endoplasmic Reticulum, (6) Golgi apparathus, (7) Cytoskeleton, (8) Smooth endoplasmic reticulum, (9) Mitochondrion, (10) Vacuole, (11) Cytoplasm, (12) Lysosome, (13) Vescicles. B) The peroxisome and its structure, showing the lipidic bilayer membrane, the inner matrix and crystalloid core (not always present). Peroxisomal proteins can be divided in two groups, matrix and membrane proteins, depending on the localisation. Membrane proteins are found attached on the inner and outer surface or can span through the layer (trans-membrane proteins). Panel A is partially adapted from en.wikipedia.org/wiki/Ribosome#/media/File:Animal Cell.svg.

The organelles provide suitable biological conditions for proteins and the correct transport of a protein to its final destination is crucial to its function. Failure in protein transport systems has been associated with several disorders including Alzheimer’s and cancers [2–4].

It has been observed that proteins from different organelles show signals, in their amino acid composition, specifically for their subcellular localisation [5]. This lead to the hypothesis that each protein has evolved to function optimally in a given subcellular compartment and to the idea that information encoded in the sequence can be used to predict the subcellular organisation. Since the pioneering work of Nakashima and Nishikawa, who used the amino acid composition to discriminate between intra- and extra-cellular proteins [6], several studies have been proposed to predict protein localisation (see [1] for comprehensive reviews). A list of the most common tools for subcellular localisation includes BaCello [7] a predictor based on different Support Vector Machines (SVM) organised in a decision tree; Phobius [8], a combined transmembrane topology and signal peptide predictor; WoLF PSORT [9] a *k*-nearest neighbors based classifier; TPpred3 [10], an SVM predictor exploiting N-terminal targeting peptides.

While these approaches use classical machine learning strategies like SVM to build classifiers, recent advances in artificial intelligence have seen the appearance of deep-learning classifiers of unprecedented accuracy in performing certain tasks like image or sound recognition [11, 12]

Deep learning algorithms learn how to perform classification tasks directly from text sequences or images. They use artificial neural network architectures, information processing paradigms inspired by the way biological neural systems process data, and try to simulate and exploit some properties of these. In its simple form, deep learning is an evolution of the linear Perceptron [13], an algorithm used for supervised learning of binary classifiers that is a one-layer artificial neural network.

Deep learning exploits multi-layer (or deep) artificial neural networks (hence the name “deep”) and combines multiple non-linear processing layers, using simple elements operating in parallel like the brain does. The layers are interconnected via nodes, or neurons, with each hidden layer using the output of the previous layer as its input.

Given its ability of modelling phenomena that can exhibit complex global behaviour, deep learning is in principle well suited for classification problems like the determination of the subcellular localisation of a protein, where the relationship between the input (*e*.*g*. instance the protein amino acid sequence) and the output (the protein localisation) is non-linear.

Several DL approaches have been recently proposed implementing Convolutional Neural Networks (CNN) [14] and Recurrent Neural Networks (RNNs)[15].

CNNs are fully connected networks where regularisation is applied hierarchically and has been exploited in DeepMito [16, 17] for the determination of sub-mitochondrial localisation. Recurrent Neural Networks have been applied for protein sequence analysis [18–20]: the advantage of RNN is that they can process sequences retaining information from previous positions in the sequence at any specific point. Variants of the RNNs are bidirectional RNNs (BiRNN) where two RNNs, one per each sequence directions [21] is used. Each residue is then contextualised having a process that goes from the N terminus to the C terminus and vice-versa.

Other variants are implemented in DeepLoc [22] which uses Bi-directional long shortterm memory (LSTM) RNN [23] (BiLSTM) for subcellular localisation and TargetP2.0 [24] to predicts the presence of N-terminal presequences which uses BiLSTM and a multi-attention mechanism, enabling the prediction of both the type of peptide and the position of the cleavage sites.

Standard machine learning and deep learning approaches exploit either *i*) the overall protein amino acid composition, *ii*) known target sequences, *iii*) sequence homology and/or motifs, or *iv*) a combination of information from the first three categories (hybrid methods) [1].

Protein sequences are commonly transformed to numerical representations that can be mathematically manipulated. Classically, these representations are referred to as “encodings” and can be broadly subdivided in four categories *i*) binary encoding, *ii*) encoding based on physical-chemical properties, *iii*) evolution-based encoding and, *iv*) structural encoding [25]. Examples are the one-hot encoding (1HOT) [25], the residue physical-chemical properties encoding (PROP) [26], the Position-Specific Scoring Matrix (PSSM) [27].

Recently, deep learning methods have been proposed, and applied, to extract fundamental features of a protein and to embed them into a statistical representation that is semantically rich and structurally, evolutionary, and bio-physically grounded [28]. These statistical representations are known as deep-learning embeddings (DL-embeddings) and are a multidimensional transformation of the protein sequence obtained using deep learning to extract and learn the information from the huge amount of protein sequences available in biological databases. We can take advantage of these embeddings for several tasks, especially subcellular localisation [29, 30]. Two of the most promising DL-embeddings are the Unified Representation (UniRep) embedding [28] and the Sequence-to-Vector (SeqVec) embedding [31]. In particular, UniRep [28] showed aminoacid embeddings containing meaningful physicochemically and phylogenetic clusters and proved to be efficient for distinguishing proteins from various structural classifications of proteins (SCOP) classes. SeqVec also showed similar results [31] and showed optimal performances for predicting subcellular localisation including peroxisomes.

Peroxisomes (Figure 1) are ubiquitous organelles surrounded by a single biomembrane that are relevant to many metabolic pathways such ether phospholipid biosynthesis, fatty acid beta-oxidation, bile acid synthesis, docosahexaenoic acid (DHA) synthesis, fatty acid alpha-oxidation, glyoxylate metabolism, amino acid degradation, and ROS/RNS metabolism [32]. They also have several non-metabolic functions, for example in cellular stress responses, response to pathogens and antiviral defence, as cellular signalling platforms and in healthy ageing [33]. For all these reasons they gained the appellative of “protective” organelles [33] and dysfunctions in peroxisomal proteins have been associated with metabolic disorders in humans [32, 33]. Peroxisomes were first described as microbodies in 1954 [34] and later identified as organelles 1967 [35] but the full extent of their functions is still largely unknown [36]. A fundamental step to gain information about the peroxisome roles and functions is the discovery of new peroxisomal proteins and the characterisation of their function. This led to the problem to determine the sub-peroxisomal localisation of peroxisome proteins. For instance, both membrane contact site (MCS) proteins [37] and peroxisomal transporters (PT) [38] are found on the membrane: *i*.*e*. distinguishing between proteins located on the peroxisomal membrane or in its granular matrix is thus a fundamental step for the characterization of unknown peroxisome proteins.

Despite its relevance, the problem of protein sub-peroxisomal localisation has received limited attention: as for today, the only way to retrieve information about the sub-peroxisomal localisation is to check for short conserved sequence motif known as signal motifs, or protein targeting signals (PTS). That is the case of the features implemented in ‘PeroxisomeDB’ [39]. From the PeroxisomeDB webserver (www.peroxisomedb.org), it is possible to identify PEX19BS, PTS1 and PTS2 targeting signals, given a FASTA sequence as input. More precisely, PTS1 and PTS2, identify peroxisomal matrix proteins while PEX19BS identifies peroxisomal membrane proteins.

In this study, we address the problem of predicting the sub-localisation of peroxisomal protein using a computational strategy combining protein-sequence embedding with classical machine learning. We reviewed and compared the four different machine learning approaches, namely Logistic Regression (LR), Random Forest (RF), Support Vector Machine (SVM), and Partial Least Square Discriminant Analysis (PLS-DA) in combination with five different protein embedding approaches namely residue one-hot encoding (1HOT), residue physical-chemical properties (PROP), Position Specific Scoring Matrices (PSSM), Unified Representation (UniRep), Sequence-to-Vector (SeqVec).

Based on our comparative study, we built a computational pipeline (In-Pero) which is based on Support Vector Machines and the combination of UniRep and SeqVec embedding. We also tested our approach for sub-mitochondrial localisation, obtaining a predictor that outperformed most of the existing classifiers.

## Methods

### Overview of the full comparison workflow

A complete overview of the comparison strategy for the selection of the best classification strategy to predict the sub-localisation of peroxisomal proteins is given in Figure 2. The comparison pipeline consists of three main steps:

**Figure 2.**
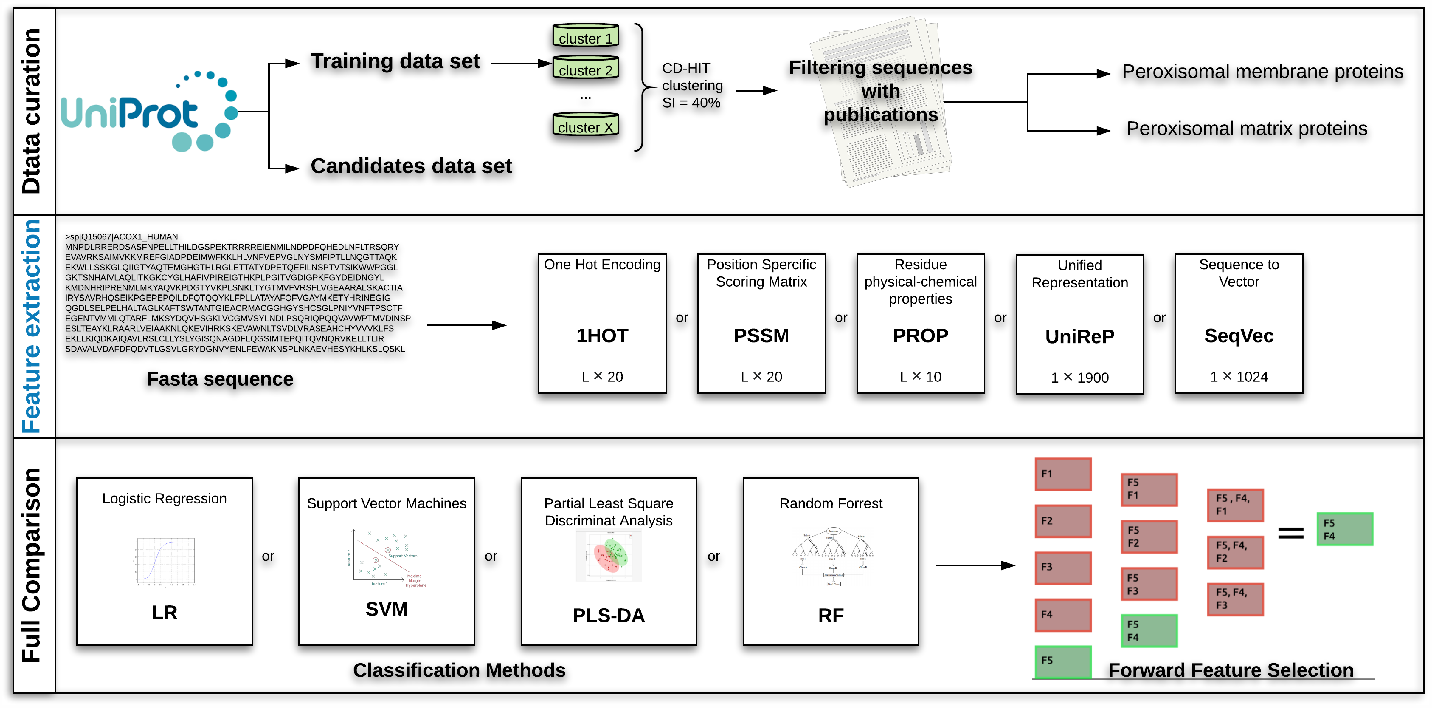
Overview of the full analysis for the predictor pipeline development. Data curation: retrieval and selection of peroxisomal protein sequences (see Sections 1). Feature extraction: conversion of protein sequences to standard encodings, namely: one-hot encoding (1HOT), residue physical-chemical properties encoding (PROP), position specific scoring matrix (PSSM), unified representation (UniRep), sequence-to-vector (SeqVec). Full Comparison: application of classification algorithms (Section 1) and selection of the best combination(s) of sequence encodings and embeddings using step forward feature selection (see Section 1)

1. Data curation: Retrieval of peroxisome protein sequence from UniProt, clustering and filtering.
2. Feature extraction: Transformation of the protein sequences into numerical representations capturing protein characteristics using classical encodings (1HOT, PROP and PSSM) and deep-learning embeddings (UniRep and SeqVec).
3. Full comparison. Double cross-validated assessment of the prediction capability of different combination of machine-learning approaches (Logistic regression, Support vector machines, Partial least square discrimination analysis and Random Forest) and protein sequence encodings and embeddings using Step Forward Feature Selection.

All methods and approaches used are detailed in the following sections.

### Data sets

Amino acid sequences for peroxisomal membrane and matrix proteins were retrieved in December 2019 from the UniprotKB/SwissProt database (www.uniprot.org) [40].

To build a highly curated data set we followed a similar approach to the one used to create data sets used to train and build predictor for the subcellular localisation of mitochondrial protein as previously reported [16, 41]. Sequences were retrieved and processed as described below.

#### Retrieval of peroxisomal membrane proteins

Peroxisomal membrane proteins were retrieved using the query ‘fragment:no locations:(location:”Peroxisome membrane [SL-0203]”) AND reviewed:yes’ with peroxisomal membrane sub-cellular location (SL-0203) to select select reviewed, non-fragmented membrane protein sequences.

We obtained 327 non-fragmented protein sequences which were then clustered using Cd-hit [42], with sequence identity of 40%. The representative (*i*.*e*. the longest protein sequence in the cluster) of each cluster was chosen resulting 162 sequences.

We restricted further the selection only to those proteins with at least one associated publication specific for the sub-cellular localization, obtaining 135 highly curated peroxisomal membrane protein sequences. Additionally, three sequences were removed from the data set, since they were not available for the UniRep embedding. The final data set contains 132 membrane proteins from four different kingdoms (see Table 1).

**Table 1.** Summary of peroxisomal protein sequences used in the present study. Sequences were retrieved and curated as detailed in Section 1.

|  | Fungi | Metazoa | Viridiplanta | Protozoa |
| --- | --- | --- | --- | --- |
| Matrix | 15 | 9 | 4 | 0 |
| Membrane | 29 | 38 | 54 | 11 |

#### Retrieval of peroxisomal matrix proteins

Peroxisomal matrix proteins were obtained with the query ‘fragment:no locations:(location :”Peroxisome matrix [SL-0202]”) AND reviewed:yes’ to select reviewed, non-fragmented matrix protein sequences. We obtained 60 entries that were further reduced to 22 after the clustering and to 19 after selecting only those proteins with at least one publication specific for the subcellular localisation.

Due to the low number of matrix proteins with comparison to the number of membrane proteins (132) we performed another advanced search in Uniprot with query ‘fragment:no locations:(location:”Peroxisome [SL-0204]”) NOT locations:(location:”Peroxisome membrane [SL-0203]”) AND reviewed:yes’ selecting reviewed, non-fragmented protein sequences, with peroxisomal location (SL-0204), and not peroxisomal membrane location (SL-0203).

We obtained 721 non membrane protein sequences, 202 after clustering, which were reduce to 22 after applying the same filtering procedure. There were 13 common entries between the two subsets; clustering using a 40% sequence similarity threshold gave 28 unique peroxisomal matrix protein sequences from four kingdoms (see Table 1).

#### Retrieval of candidate peroxisomal proteins

Further peroxisomal protein candidates were retrieved from Uniprot (Jun 2020). We looked for peroxisomal proteins (SL-0204 and GO:5777) with a non-specific sub-peroxisomal location (SL-0203, SL-0202, GO:5778, GO:5782) and experimental evidence. We then excluded the peroxisomal proteins also found in mitochondria (SL-0173, GO:5739) and in the endoplasmic reticulum (SL-0095, GO:10168) obtaining 116 reviewed entries.

#### Data sets for sub-mitochondrial protein classification

To assess the generalizability of our prediction tool to the prediction of the subcellular localisation of protein from other organelles we considered two well-curated data sets containing mitochondrial proteins

##### SM424-18 data set

this data set was used to build the DeepMito predictor [16] and contains 424 mitochondrial proteins collected using stringent conditions, in particular only non-fragmented proteins with and experimentally determined subcellular localisation in one of the four sub-mitochondrial compartments (outer membrane, inter-membrane space, inner membrane and matrix). Clustering using Cd-hit [42], with a 40% sequence identity threshold was used to select representative sequences. We refer the reader to the original publication for more details [16].

##### SubMitoPred data set

this data set was used to build the SubMitoPred predictor [41]. It contains 570 mitochondrial proteins collected using stringent conditions, in particular only non-fragmented proteins with and experimentally determined subcellular localisation in one of the four sub-mitochondrial compartments (outer membrane, inter-membrane space, inner membrane and matrix). Clustering using Cd-hit [42], with a 40% sequence identity threshold was used to select representative sequences. We refer the reader to the original publication for more details [41].

### Classic protein sequence encoding methods

We considered three of the most commonly used method for the encoding of the amino acid protein sequences.

#### Residue one-hot encoding

The one-hot encoding (1-HOT) [25] is the most used binary encoding method. Any given residue *j* is represented by a 1 *×* 20 vector containing 0s except in the *j*-th position; for instance alanine (A) is represented as 100000000000000000000. A protein sequence constituted by *L* amino acid is thus represented by an *L ×* 20 matrix.

#### Residue physical-chemical properties encoding

Akinori *et al*. devised a way to represent an amino-acid with ten factors [26] summarising different amino acid physico-chemical properties. This encoding method, often abbreviated as PROP, is the most commonly used physico-chemical encoding [25].

Any given residue *j* in the protein sequence is represented by a 1 *×* 10 vector containing real number. Each number summarise different amino-acid properties and it is an orthogonal property obtained after multivariate statistical analysis applied to a starting set of 188 residue-specific physical properties. A protein sequence constituted by *L* amino acid is thus represented by an *L ×* 10 matrix.

#### Position specific scoring matrices

The Position-specific scoring matrix (PSSM) [27] takes into account the evolutionary information of a protein. This scoring matrix is at the basis of protein BLAST searches (BLAST and PSI-BLAST) [43] where residues are translated into substitution scores.

Any given residue *j* in the protein sequence is represented by a 1 *×* 20 vector containing the 20 specific substitution scores. Each amino acid substitution scores are given separately for each position in a protein multiple sequence alignment (MSA) after running PSI-BLAST [43] against the Uniref90 data set (release Oct 2019) for three iterations and e-value threshold set to 0.001. We used a sigmoid function to map the values extracted from the PSI-BLAST checkpoint file in the range [0-1], as in DeepMito [16]. Basically, PSSM captures the conservation pattern in the alignment and summarises evolutionary information of the protein.

In PSSM a protein sequence constituted by *L* amino acid is thus represented by an *L ×* 20 matrix.

### Deep learning protein sequence embeddings

We considered two recently proposed methods for the embedding of protein sequences based on deep-learning approaches

#### Unified Representation

The Unified Representation (UniRep) [28] is based on a recurrent neural network architecture (1900-hidden unite) able to learn statistical representations of proteins starting from *∼* 24 million UniRef50 sequences.

UniRep is based on multiplicative long-/short-term-memory (mLSTM) RNNs [44] which is a combination of LSTM, which uses multiplicative gates to control how information flows trough internal states of the network [45], and mRNN [20] is designed to allow flexible input-dependent transitions.

Technically, the protein sequence is modelled by using a hidden state vector which is recursively updated based on the previous hidden state vector. This means that the method learns scanning a sequence of amino acids, predicting the next one based on the sequence it has seen so far.

The UniRep statistical has been proved able to capture chemical, biological and evolutionary information encoded in the protein sequence and has been used to obtain cluster of proteins according to physico-chemical properties, organism origin at different phylogenetic levels and to SCOP (structural classifications of proteins) classification [46]

Using UniRep a protein sequence can be represented by an embedding of length 64, 256, or 1900 units depending on the neural network architecture used. In this study, we used the 1900 units long (average final hidden array).

#### Sequence-to-Vector embedding

The Sequence-to-Vector embedding (SeqVec) [31] embeds biophysical information of a protein sequence taking a natural language processing approach considering amino acids as words and proteins as sentences.

SeqVec is obtained by training ELMo [47], a deep contextualised word representation that models both complex characteristics of word use (e.g., syntax and semantics), and how these uses vary across linguistic contexts, which consists in a 2-layer bidirectional LSTM backbone pre-trained on a large text corpus, in this case, UniReg50.

The SeqVec embedding can be obtained by training ELMo at the per-residue (word-level) and per-protein (sentence-level). With the per-residue level it is possible to obtain a protein sequence embedding that can be use to predict the secondary structure or intrinsically disordered region; with the per-protein level embedding it is possible to predict subcellular localisation and to distinguish membrane-bound vs water-soluble proteins [31]. Here we use the per-protein level representation, where the protein sequence is represented by an embedding of length 1024.

### Step Forward Feature Selection

Step Forward Feature Selection was used to select the best combination of features (predictors) *i*.*e*. protein encodings or embeddings to be used as input for classification algorithms [48].

It is a wrapper method that evaluates subsets of variables, in our case combinations of protein encodings/embeddings. It starts with the evaluation of each individual encoding, and selects that which results in the best performing selected algorithm model. Next, it proceeds by iteratively adding one encoding/embedding to the current best performing features and evaluating the performance of the classification. The procedure is halted when performance worsens and the best combination of embeddings/encodings is retained. A schematic representation of this approach given in Figure 2 (Step 3: Full Comparison).

### Classification algorithms

The determination of the sub-localisation of peroxisomal (membrane *vs* matrix) protein is easily translated into a two-group classification problem. For this we considered four widely used machine learning methods which are described in the following section with the relevant meta-parameters.

### Support Vector Machines

Support Vector Machines (SVM) is an algorithm for two-group classification which aims to find the maximal margin hyperplane separating the points in the feature space [49, 50]. The SVM algorithm has three hyperparameters that require optimization:

1. The cost parameter *C* that controls the trade off between errors of the SVM on training data and margin maximization; basically *C* that controls the missclassifcition.
2. The type of kernel, *i*.*e*. the type of transformation applied to the input data like linear, nonlinear, polynomial, radial basis function (RBF), and sigmoid.
3. *γ* a specific parameter for the radial basis kernel.

### Random Forest

Random Forest [51, 52] is an ensemble learning method that, in the case of a classification task construct a multitude of decision trees and output the mode of the classes of the individual trees. The RF algorithm has several hyperparameters that require optimisation. Using the implementation available in scikit-learn [53], we optimised the following parameters:

1. ‘n_estimators’ the number of trees in the forest
2. ‘criterion’ the function to measure the quality of a split.
3. ‘max_depth’ the maximum depth of the tree.
4. ‘min_samples split’ the minimum number of samples required to split an internal node.
5. ‘max_features’ the number of variables to consider when looking for the best split.

### Partial Least Square - Discriminant Analysis

Partial least squares discriminant analysis (PLS-DA) is a partial least squares regression [54, 55] where the response vector *Y* contains dummy variables indicating class labels (0-1 in this case). Sample predicted with *Y ≥* 0.5 are classified as belonging to class 1 and to class 0 other wise. PLS finds combinations of the original variable maximizing the covariance between the predictor variable and response *Y* by projecting the data in a *k*-dimensional space with *k* possibly much smaller than the original number of variables. The PLS algorithm has only one hyperparameter: the number of number of components *k*.

### Logistic regression

We used a penalised implementation of multivariable logistic regression [56] available in scikit-learn [53] which has three metaparamters to be tuned:

1. ‘solver’ the algorithm (method) to use in the optimisation problem (e.g.’liblinear’, ‘saga’, etc…)
2. ‘penalty’ the norm used for penalisation *L*_1_ or *L*_2_
3. The cost parameter *C* controlling the bias between model fit and miss-classification.

## Model calibration and validation

We used double cross validation (DCV) [57, 58] for *i*) optimising the hyper-parameters of the different classification algorithm used (*i*.*e*. for model calibration) and *ii*) for an unbiased estimation of prediction errors when the model is applied to new cases (that are within the population of the data used). This strategy is particularly well suited for small data sets.

The DCV strategy consists in two nested cross-validation loops. In the outer loop data is first split in *k* folds. One fold is used as Validation set while the remaining *k −* 1 folds are used as calibration set. The inner loop is applied to the Calibration set which is again split in a test and training set using *k*-fold split. The inner loop is used to optimise the hyperparameters of the different classification algorithms through a (hyper)grid search: for each set of hyperparameters, the average classification score is computed across the folds. The hyperparameters corresponding to the best classification score are then used to fit a classification model whose quality is assessed on the Validation set obtaining unbiased model evaluation since the validation data has not been used to train the classification model.

The step forward feature selection procedure described in Section 1 was included in the calibration loop so that model calibration involved also the selection of the best combination (with respect with model predictive ability) of protein sequence encodings and embeddings.

Given the unbalance of the two class of proteins, different weights were applied to the two class. Class weights were considered as metaparameters and optimised in the inner calibration loop.

### Metrics for model classification accuracy

We used several metrics to quantify the quality of the classification models.Accuracy (*ACC*) *i*.*e*. the classification error, defined as

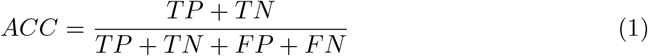

where: *TP* is the number of true positives, *FP* is the number of false positives; *TN* and *FN* are the number of true and false negatives, respectively.

The *F*_1_ score [59]:

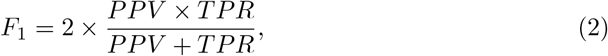

where *PPV* is the as positive predicted value (or precision)

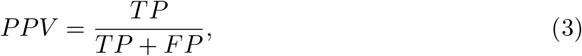

and *TPR* is the true positive rate (recall or sensitivity):

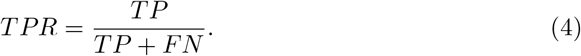

The *F*_1_ score is the harmonic mean of recall and precision and varies between 0, if the precision or the recall is 0, and 1 indicating perfect precision and recall.

The balanced accuracy *BACC* [60]:

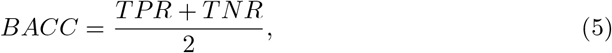

where

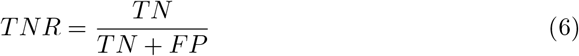

is the true negative rate or specificity. The *BACC* is an appropriate measure when data is unbalanced and there is no preference for the accurate prediction of one of the two classes.

The Matthews correlation coefficient (MCC) [61]:

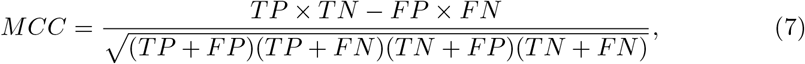

*MCC* is the correlation coefficient between the true ad predicted class: it is bound between −1 (total disagreement between prediction and observation) and +1 (perfect prediction); 0 indicates no better than random prediction. The *MCC* is appropriate also in presence of class unbalance [62].

### Prediction of trans-membrane proteins

We used TMHMM (Trans-Membrane Hidden Markov Model)[63, 64] for the prediction of trans-membrane proteins.

TMHMM returns the most probable location and orientation of trans-membrane helices in the protein sequence, summarised in several output parameters: ExpAA, the expected number of amino acids in transmembrane helices. If this number is larger than 18 it is very likely to be a transmembrane protein; First60, the expected number of amino acids in transmembrane helices in the first 60 amino acids of the protein; PredHel, the number of predicted transmembrane helices. We refer the reader to the original publications for more details.

We used TMHMM to predict the localisation of peroxisomal protein with unknown sub-localisation (see data sets description in Sections *Retrieval of candidate peroxisomal* proteins)

### Software

For all classification algorithms we used the implementation available in the scikit-learn python library (version 0.22.1) [53]. We obtained the PLS-DA algorithm by adapting the PLS regression algorithm to perform a regression with a dummy variable.

## Results

### Selection of the best classifier for sub-peroxisomal prediction

We compared four commonly used machine learning approaches (Logistic regression, Partial Least Squares Discriminant analysis, Random Forest and Support Vector Machines) in combination with different protein sequence encodings and embeddings to select the best classification strategy to predict the sub-localisation of peroxisomal proteins. Results are summarised, per classification algorithm, in Table 2; where different metrics for model quality quantification are given. All results were obtained with repeated double cross-validation to avoid model overfitting and bias.

**Table 2.**
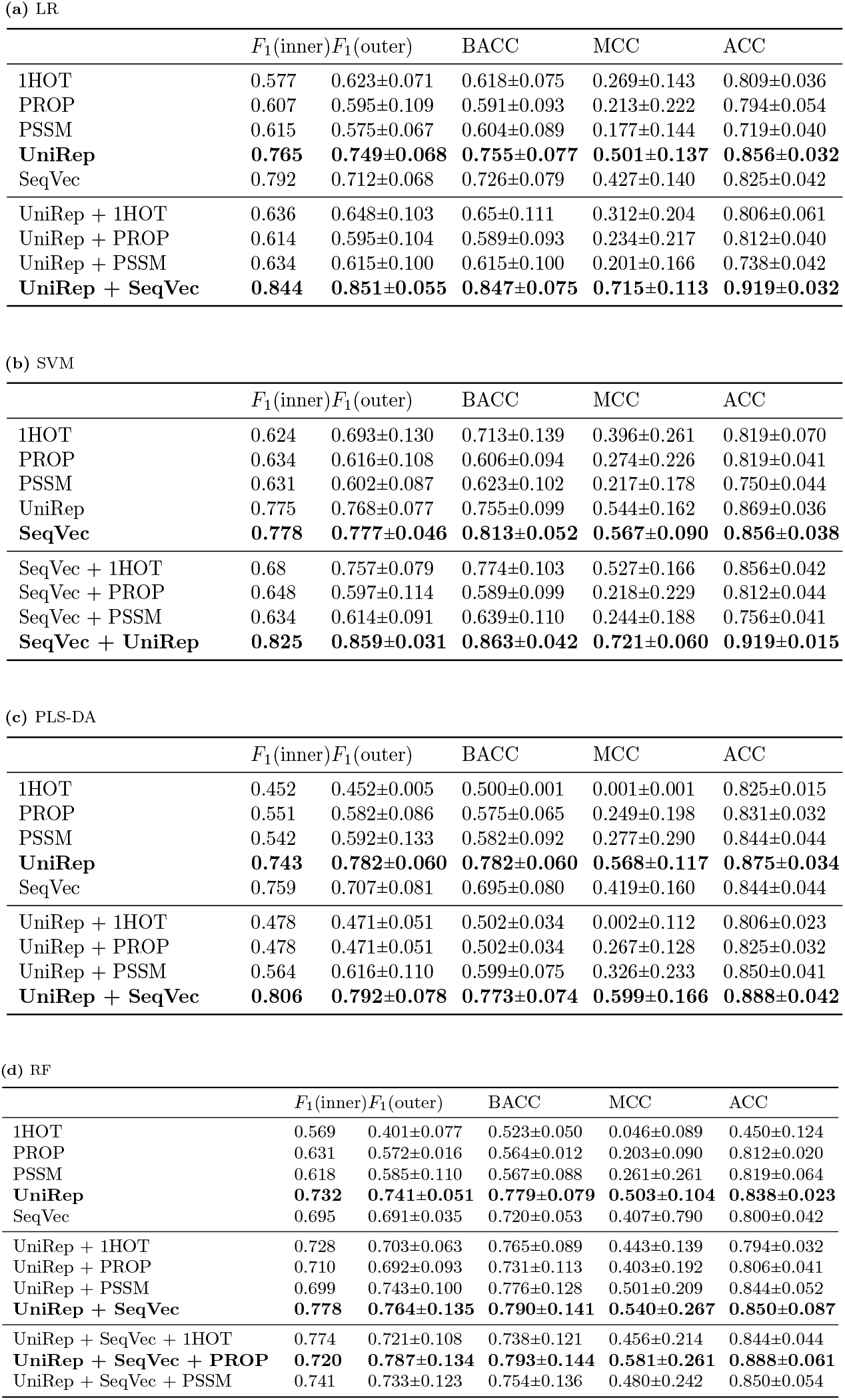
Step Forward Feature Selection for each of the compared methods. a) Logistic Regression (LR) performances. b) Support Vector Machines (SVM) performances. c) Partial Least Square - Discriminant Analysis (PLS-DA) performances. d) Random Forest (RF) performances. The analysed encodings and embeddings are protein one hot encoding (1HOT), residue physical-chemical properties encoding (PROP), position specific scoring matrix (PSSM), Unified Representation (UniRep) and Sequence to Vector (SeqVec). The results refer to the Double Cross Validation (DCV) procedure performed for each iteration of the forward feature selection (Section *Step Forward Feature Felection*). The *F*_1_ (inner) score refers to the inner loop of the DCV while *F*_1_ (outer) refers to the outer loop. The performances are reported in terms of F1 score (equation 2), BACC (equation 5), MCC (equation 7) and ACC (equation 1).

In general, Logistic regression (Table 2a) and Support vector machines (Table 2b) showed similar performance superior to PLS-DA 2c) and Random Forest (Table 2d). However, in general, the prediction model built using SVM has a smaller standard deviation, indication larger model stability.

We observed that combining two different encodings and/or embeddings gives a better prediction of the peroxisomal sub-localisation. In particular, concatenating UniRep and SeqVec showed a noticeable improvement in the performances. That indicates the two embeddings carry different and complementary information about the properties of the protein sequence. As shown in Figure 3, the two embeddings are not correlated.

**Figure 3.**
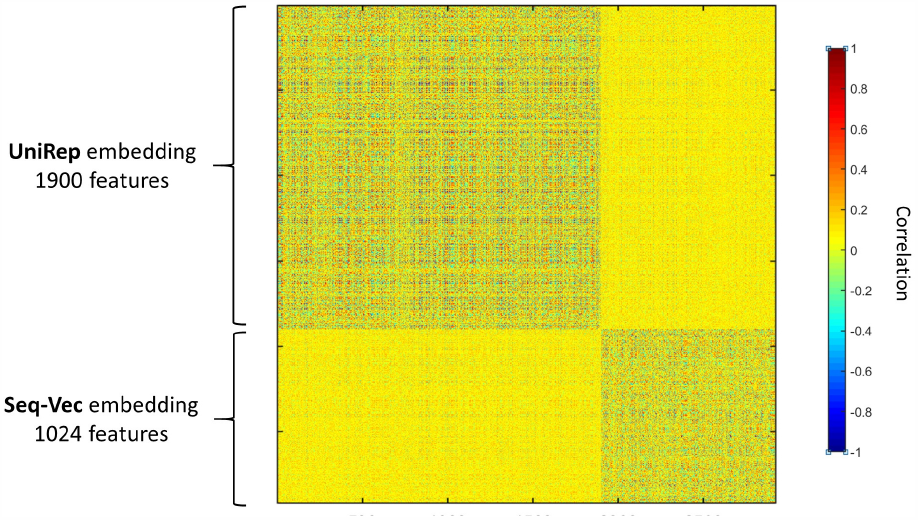
Correlation among the UniRep (1900 features) and the SeqVec (1024 features) protein sequence embeddings. Pearson’s linear correlation is used and are calculated over 160 protein sequences. The two embeddings are uncorrelated.

### In-PERO a tool for the prediction of peroxisomal protein sub-localisation

Based on the results obtained and discussed in Section *Selection of the best classifier for sub-peroxisomal prediction* we developed In-Pero, a computational pipeline to predict the Intra-Peroxisomal localisation of a proximal protein, *i*.*e*. to discriminate between matrix and membrane proteins. In-Pero relies on Support Vector Machine classifier trained on the statistical representation of protein sequences obtained by the combination of two deep-learning embeddings.

In-Pero consists of four main steps (See Figure 4A)

**Figure 4.**
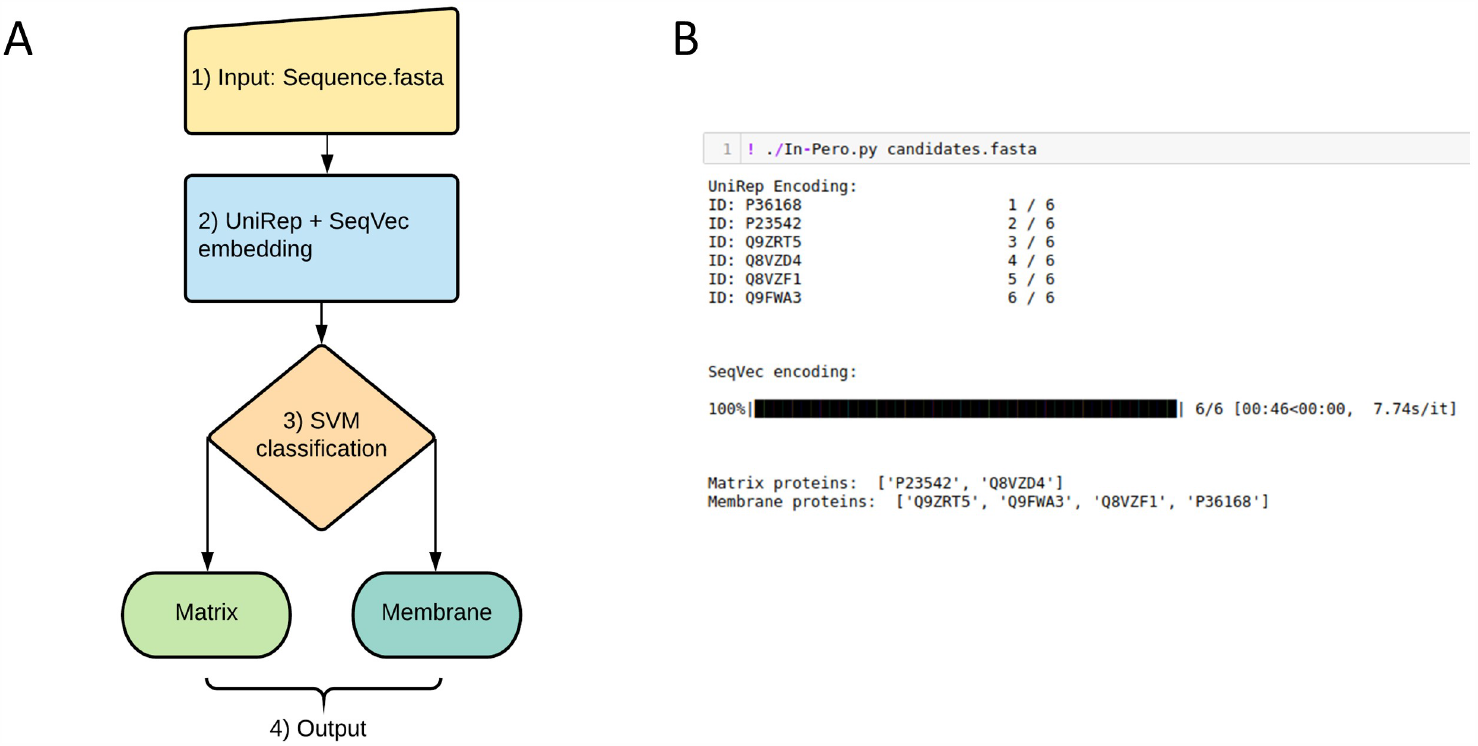
A) In-Pero workflow 1) Input as protein fasta sequence. 2) Sequence representation via DL encoding, in particular concatenating UniRep and SeqVec. 3) Support Vector Machines based classification. 4) Output the sub-peroxisomal location. B) Example of a typical execution, with 6 sequences contained in the candidates.fasta file: the sub-peroxisomal classification of each protein is give.

1. Input of the protein sequence in FASTA format.
2. Calculation of the statistical representation of the protein sequence using the UniRep (1 *×* 1900) and the SeqVec (1 *×* 1024) embeddings
3. Merging of the two statistical representation to obtain a 2924-dimensional representation of the protein sequence.
4. Prediction of the sub-cellular localization using the trained SVM

In-Pero is implemented in Python and work in command line modality. An example of the input command line and output is given in 4B). A link to the In-Pero code is available in the Availability Section and in the Abstract).

### Validation of sub-peroxisomal membrane protein prediction

The In-Pero prediction tool was trained and validated using a double cross-validation strategy (see Section *Model calibration and validation*) with a stratified 5-fold splitting. The predictive capability of the model was assessed on the data that have not been used for model calibration (*i*.*e*. the selection of meta-parameters giving the best prediction quality). This approach is a proxy for the use of an external data set for experimental validation, ensures unbiased model assessment and reduce the risk of over-fitting.

Despite all precautions, it is important to benchmark In-Pero against existing tools. There are no existing computational tools for the sub-localisation of peroxisomal protein. As a work-around, we compared the prediction of In-Pero with those of TMHMM using a set of 116 peroxisomal protein of unknown sub-peroxisomal localisation (see Section *Retrieval of candidate peroxisomal proteins*) which have not been used to train the In-Pero classifier.

When In-Pero is run on these 116 proteins we obtained membrane localisation for 48 and matrix localisation for 68. We tested the 48 protein classified as membrane proteins using TMHMM: 7 were predicted as transmembrane proteins while 13 have characteristics compatible with transmembrane localisation (a value *>*= 1 for at least one among the ExpAA, First60 and PredHel scores, see Section *Prediction of trans-membrane* proteins). The results are visible in Table 3.

**Table 3.** Transmembrane and Membrane proteins found with both methods (In-Pero and TMHMM). The seven Transmembrane proteins showing high prediction scores with TMHMM are in bold.

| Membrane |  | Transmembrane |
| --- | --- | --- |
| O82399 | Q8H191 | **O64883** |
| P36168 | Q8K459 | **P20138** |
| P90551 | Q8VZF1 | **Q84P23** |
| Q12524 | Q9LYT1 | **Q84P17** |
| Q4WR83 | Q9NKW1 | **Q84P21** |
| Q75LJ4 | Q9S9W2 | **Q9M0X9** |
| Q9SKX5 |  | **P08659** |

Among the seven classified as transmembrane proteins, two (064883, P20138) showed experimental evidence in being connected to various cellular membranes while the sub-peroxisomal location is not reported. Among the others, four (Q84P23, Q84P17, Q84P21, Q9M0X9) are present in *Arabidopsis thaliana* while P08659 is present in *Photinus pyralis*. For these five entries the membrane association is not reported, neither the sub-peroxisomal location, making them candidates for a more precise annotation.

It should be noted that TMHMM method focuses on predicting transmembrane regions, not predicting subcellular location. However, we can speculate that also peroxisome membrane protein may share some structural and physico-chemical properties similar to cell membrane proteins; thus confronting the results of TMHMM with In-Pero can provide partial, independent validation of the In-Pero classifier.

### Extending In-Pero to predict Sub-mitochondrial proteins

To explore further the range of applicability of the combination of machine learning and deep-learning protein sequence embeddings to other problems related to the prediction of protein localisation, we applied In-Pero for sub-mitochondrial classification. Mitochondrial proteins are physiologically active in different compartments (the matrix, the internal membrane, the inter-membrane space and the external membrane) and their aberrant localisation contributes to the pathogenesis of human mitochondrial pathologies [65]. By adapting In-Pero to a multiclass classification problem, we obtained the In-Mito predictor. We considered both an SVM and a Logistic regression as classification algorithm since, in this case, they performed similarly.

There are several tools available for the prediction of sub-mitochondrial localisation. We compared In-Mito against SubMitoPred [41], DeepMito [16], and DeepPred-SubMito [66].

We tested our model with the SM424-18 and SubMitoPred data sets (see Section *Data sets for sub-mitochondrial protein classification*): results are given in Table 4.

**Table 4.** Comparison with DeepMito and DeepPred-SubMito (DP-SM) based on the SM424-18 data set (Data A) and the SubMitoPred data set (Data B). The results are reported in terms of Mattew Correlation Coefficient (MCC) explained in Equation The four mitochondrial compartments are outer membrane (O), inner membrane (I), intermembrane space (T) and matrix (M). Given the similar performances of our predictor (In-Mito) implemented with Logistic Regression (LR) and Support Vector Machines (SVM), we report both. The best performances are in bold.

| Data A | MCC(O) | MCC(I) | MCC(T) | MCC(M) |
| --- | --- | --- | --- | --- |
| DeepMito | 0.460 | 0.470 | 0.530 | 0.650 |
| DP-SM | **0.850** | 0.490 | **0.990** | 0.560 |
| In-Mito (LR) | 0.680 | **0.730** | 0.690 | **0.820** |
| In-Mito (SVM) | 0.640 | 0.690 | 0.620 | 0.800 |

| Data B | MCC(O) | MCC(I) | MCC(T) | MCC(M) |
| --- | --- | --- | --- | --- |
| SubMitoPred | 0.420 | 0.340 | 0.190 | 0.510 |
| DeepMito | 0.450 | 0.680 | 0.540 | 0.790 |
| DP-SM | **0.920** | 0.690 | **0.970** | 0.730 |
| In-Mito (LR) | 0.690 | 0.750 | 0.620 | **0.850** |
| In-Mito (SVM) | 0.650 | **0.760** | 0.540 | 0.840 |

In-Mito compared favourably to existing predictors, principally with regard to the few designed to classify all four mitochondrial compartments. Moreover, it shows a balanced capability to predict different compartments. In particular, In-Mito shows excellent performance in the prediction of matrix proteins and of inner membrane proteins, which are the most abundant subcellular compartments (80% of the SubMitoPred data set).

We note that, for this multi-class problem, we obtained better prediction performance of different sub-cellular localisation using either logistic regression (for matrix protein) or SVM (for inter-membrane proteins). This supports the idea of possibly combining different predictors for better classification.

Given the accuracy of the classifications obtained with our approach, we implemented the tool In-Mito for sub-mitochondrial classification, which works in the same way as In-Pero (Fig 4) but specifically for mitochondria. In particular, the final output here consists of one among the four possible sub-mitochondrial compartments.

## 2 Conclusions

With this work, we covered a less explored area of bioinformatics and protein sequences analysis, namely the computational prediction of the localisation of peroxisome proteins.

Building on existing approaches, we addressed the problem by combining machine learning algorithm with different combinations of protein encodings and embeddings.

We found that the (combination of) deep learning embeddings (Seq-Vec [31] and UniRep [28] outperformed classical encodings when applied to sub-peroxisomal classification. Our newly proposed prediction tool In-Pero obtained a (double cross-validated) classification accuracy of 0.92.

We further adopted the approach deployed In-Pero to predicting the subcellular localisation of mitochondrial proteins, resulting in the In-Mito classifier. We found In-Mito to compare favourably with state-of-the-art approaches and for certain classes of proteins (matrix and intermembrane) to outperform existing prediction tools like DeepMito [17] and SubMitoPred [41].

These results suggest that *i*) the evolutionary, biochemical and structural information encoded in a protein amino acid sequence cannot be fully captured by one single embedding and that different approaches need to be combined, *ii*) deep-learning embeddings are highly versatile and could become a standard for protein sequence representation and analysis and *iii*) the possibility of extending In-Pero and In-Mito for the characterisation of other sub-organelles proteins.

Moreover, while in this work we utilised machine learning approaches, we anticipate that our method can be extended to the use of deep-learning methods also for the prediction, such as convolutional neural networks, recurrent neural networks or a combination thereof.

The lack of predictors and tools specifically dedicated to the prediction of sub-localisation of peroxisomal protein makes our work the very first on this subject and presents a complete method and features benchmark that can be used as a base for future studies.

## Availability

All presented tools and data are free to use and available online. The data sets used in this study are available at https://github.com/MarcoAnteghini/In-Pero/tree/master/Dataset. Standalone versions of In-Pero and In-Mito are available at https://github.com/MarcoAnteghini/In-Pero and https://github.com/MarcoAnteghini/In-Mito.

The data set containing the 116 investigated proteins as candidates and the related prediction is available for experimental classification and proper UniProt annotations at https://github.com/MarcoAnteghini/In-Pero/tree/master/Candidates.

## Acknowledgements

We thank Katarina Elez (FU Berlin) for the in-depth discussion about deep-learning encoding procedures.

